# Axe: rapid, competitive sequence read demultiplexing using a trie

**DOI:** 10.1101/160606

**Authors:** Kevin D. Murray, Justin O. Borevitz

## Abstract

Here we implement a rapid algorithm for demultiplexing DNA sequence reads with in-read indices. Axe selects the optimal index present in a sequence read, even in the presence of sequencing errors. The algorithm is able to handle combinatorial indexing, indices of differing length, and several mismatches per index sequence.

## 1 Introduction

The incredible yield of modern DNA sequencing technologies has enabled the multiplexing of DNA samples into a single sequencing unit. Multiplexing is achieved by the addition of short sequences (indices) to each molecule to be sequenced. When sequenced, these index sequences uniquely identify the sample to which a sequence read belongs.

Many commercial protocols use platform specific features to add these DNA indices such that sequencing platforms can automatically demultiplex these samples. However, many custom sequencing protocols, including GBS [2], add indices which end users must themselves demultiplex. Combinatorial indexing schemes add independent index sequences to both pairs of a paired-end sequencing protocol, and samples are identified by the combination of these two index sequences.

Many sequencing read demultiplexers have been published. For example, both Flexbar [1] and the Fastxtoolkit’s `fastx_barcode_splitter.pl` [3] accept single- and paired-end reads, however they cannot demultiplex combinatorial indices. AdapterRemoval [6] can demultiplex combinatorial indices, but cannot demultiplex indexes which differ in length. We developed axe to address these shortcomings.

## 2 Methods

### 2.1 Implementation

`Axe` matches the prefix of a sequence read against a pre-computed trie of index sequences. To do so, `axe` first calculates all sequences within a given hamming distance of each index sequence. Axe then associates each of these sequences with its respective sample identifier using a double-array trie. This data structure allows rapid lookup of variable length prefixes within a set of index sequences, an 𝒪(1) operation with respect to the number of indices.

Reads are demultiplexed by finding the read’s longest prefix, in the trie of index sequences, and assigning that read to its associated sample. This algorithm extends easily to combinatorial indexing, where two independent indices prefix each read of a read pair. Although this algorithm is agnostic as to which end of a sequencing read contains a index, only 5’ (prefix) index demultiplexing is currently implemented.

### 2.2 Operation

To demultiplex sequence reads, one uses the command `axe-demux`. This command takes input reads as FASTQ or FASTA files which may contain single- or paired-end reads. Paired-end reads may be interleaved, and output reads can be written in any of these formats.

Axe is implemented in the C language, available at https://github.com/kdmurray91/axe. It may be built from source code on any modern POSIX operating system (including GNU/Linux and Mac OS X). The only dependencies not bundled with the source distribution are `CMake` and `zlib`. It is available in the Debian and Ubuntu GNU/Linux distributions as the `axe-demultiplexer` software package.

### 2.3 Validation experiments

To quantify the performance of `axe` relative to similar tools, 10 million 100bp paired end reads were simulated from a random 1Mbp genome using Mason2 [4]. Sets of index sequences of various sizes (see results) were drawn from the set of all 8-mers with a minimum hamming distance of 3. Sample frequencies were drawn from a gamma distribution with a shape parameter of 2; read pairs are randomly assigned a sample from these sample frequencies. Index sequences are inserted into the 5’ end of sequences and errors added with a frequency of 10^−2.5^ (PHRED quality of 25). Combinatorial index sets were generated using the same process for each read.

These datasets were used to benchmark all operational modes of `axe`, alongside previous read demultiplexing software `flexbar`, `fastx` and `AdapterRemoval`. The precise versions and parameters for these programs, and the workflow which performs the simulations reported here, are available at https://github.com/kdmurray91/axe-experiments.

## 3 Results

### 3.1 Demultiplexing accuracy

We benchmark the speed and accuracy of `axe`, `flexbar`, `AdapterRemoval` and `fastx_barcode_splitter.pl` (hereafter “`fastx`”). When demultiplexing read pairs with an index sequence on one read only (single-end), both `axe` and `fastx` are able to perfectly demultiplex all reads, with no error and with no reads left unassigned. `AdapterRemoval` fails to assign a minuscule proportion of reads, while `flexbar` mis-assigns several percent of reads (Figure 1). When demultiplexing combinatorially indexed read pairs, `axe` again demultiplexes all reads perfectly, and `AdapterRemoval` fails to assign a small proportion. When demultiplexing reads with variable-length index sequences, `axe` performs perfectly, while `flexbar` mis-assigns several percent of reads. In all cases, `axe` is the fastest demultiplexer tested. `AdapterRemoval` performs several times slower than `axe`. `fastx` and `flexbar` perform hundreds of times slower than `axe` and `AdapterRemoval` (Figure 2).

**Figure 1.**
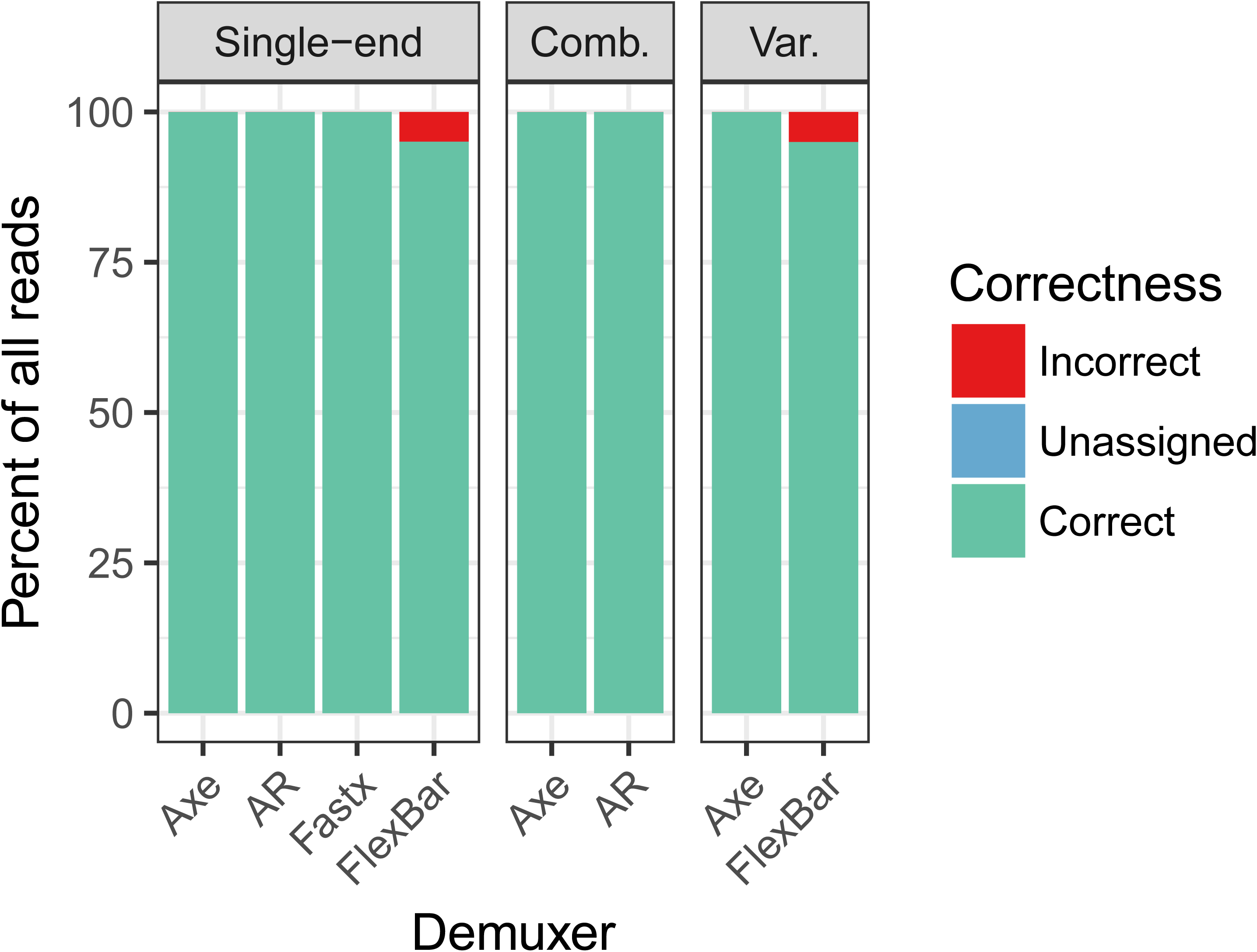
Accuracy of read assignment. `Axe` is able to perfectly demultiplex all reads, as is `fastx`. Only `flexbar` incorrectly assigns reads. Note: “Comb.” refers to combinatorial index sets, and “Var.” refers to index sets with variable length index sequences.

**Figure 2.**
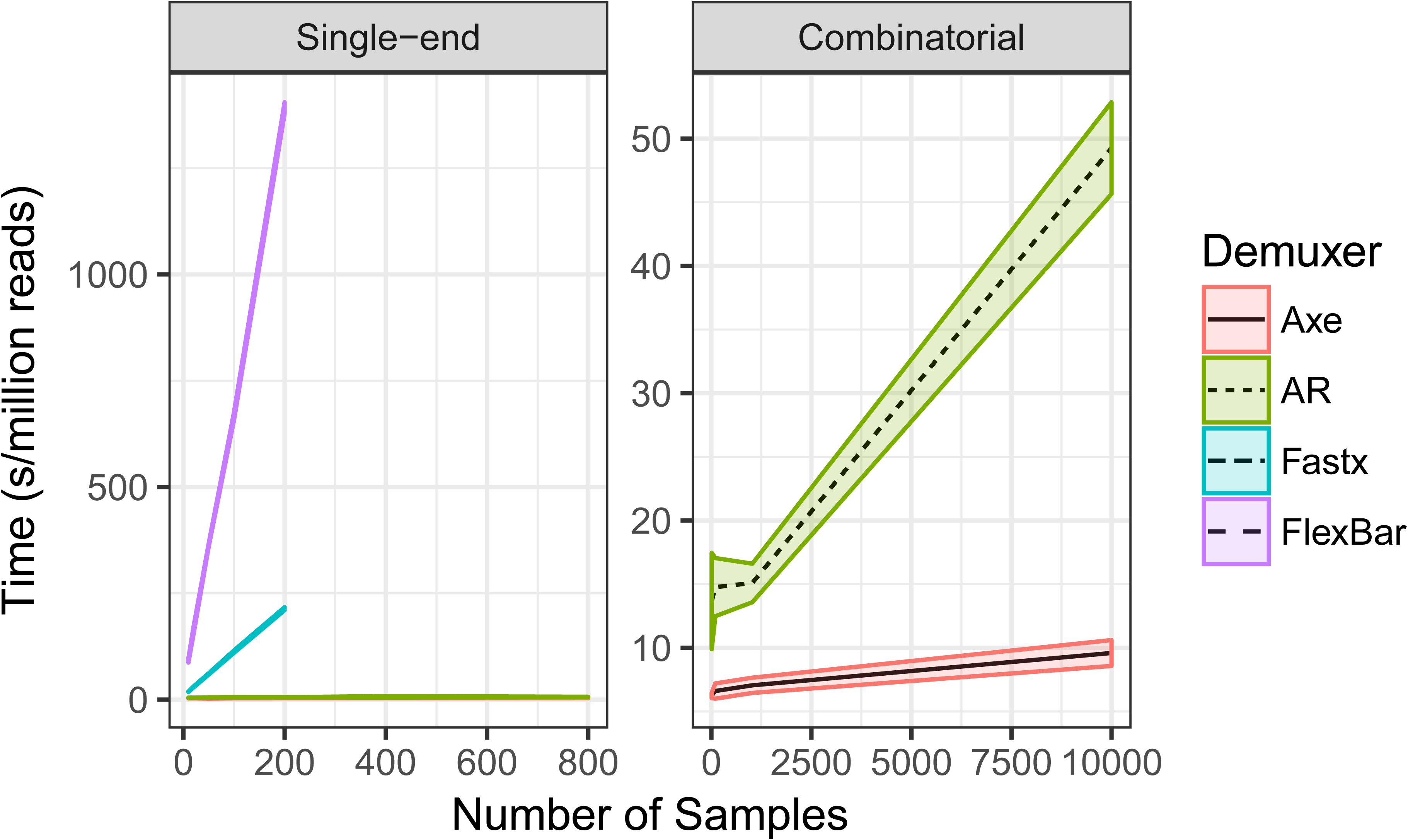
Computational performance of demultiplexers. `Axe` is the fastest in all cases, closely followed by `AdapterRemoval`. `fastx` and `flexbar` are appreciably slower, especially when the number of indices is large.

## 4 Discussion

Here we implement a rapid and accurate algorithm for demultiplexing 5’-indexed reads. We show equal or improved accuracy and reduced computational cost compared to previous software developed to perform this task. In addition, more complex indexing schemes including combinatorial and/or variable length index sequences are supported.

While in-read indexing is being phased out in some studies, it persists in protocols such as GBS[2] and RNAseq using unique molecular identifiers[5]. Additionally, `axe`’s algorithm is applicable to demultiplexing out-of-read indexing schemes, though the implementation does not currently support this.

## 5 Software Availability

### 5.1 Software available from

- Debian and Ubuntu GNU/Linux: available in the official software repositories, use `sudo apt-get install axe-demultiplexer` (for Debian/Ubuntu versions 9.0 and 16.04 or higher respectively).
- Other GNU/Linux: pre-compiled executables available from https://github.com/kdmurray91/axe/releases
- Other operating systems: portable source available from https://github.com/kdmurray91/axe/releases

### 5.2 Link to source code

https://github.com/kdmurray91/axe

## 6 Software License

Copyright 2014-2017 Kevin Murray

This program is free software: you can redistribute it and/or modify it under the terms of the GNU General Public License as published by the Free Software Foundation, either version 3 of the License, or (at your option) any later version.

This program is distributed in the hope that it will be useful, but WITHOUT ANY WARRANTY; without even the implied warranty of MERCHANTABILITY or FITNESS FOR A PARTICULAR PURPOSE. See the GNU General Public License for more details.

You should have received a copy of the GNU General Public License along with this program. If not, see http://www.gnu.org/licenses/.

